# A large-scale analysis of bioinformatics code on GitHub

**DOI:** 10.1101/321919

**Authors:** Pamela H Russell, Rachel L Johnson, Shreyas Ananthan, Benjamin Harnke, Nichole E Carlson

## Abstract

In recent years, the explosion of genomic data and bioinformatic tools has been accompanied by a growing conversation around reproducibility of results and usability of software. However, the actual state of the body of bioinformatics software remains largely unknown. The purpose of this paper is to investigate the state of source code in the bioinformatics community, specifically looking at relationships between code properties, development activity, developer communities, and software impact. To investigate these issues, we curated a list of 1,720 bioinformatics repositories on GitHub through their mention in peer-reviewed bioinformatics articles. Additionally, we included 23 high-profile repositories identified by their popularity in an online bioinformatics forum. We analyzed repository metadata, source code, development activity, and team dynamics using data made available publicly through the GitHub API, as well as article metadata. We found key relationships within our dataset, including: certain scientific topics are associated with more active code development and higher community interest in the repository; most of the code in the main dataset is written in dynamically typed languages, while most of the code in the high-profile set is statically typed; developer team size is associated with community engagement and high-profile repositories have larger teams; the proportion of female contributors decreases for high-profile repositories and with seniority level in author lists; and, multiple measures of project impact are associated with the simple variable of whether the code was modified at all after paper publication. In addition to providing the first large-scale analysis of bioinformatics code to our knowledge, our work will enable future analysis through publicly available data, code, and methods. Code to generate the dataset and reproduce the analysis is provided under the MIT license at https://github.com/pamelarussell/githubbioinformatics. Data are available at https://doi.org/10.17605/OSF.IO/UWHX8.

**Author summary:** We present, to our knowledge, the first large-scale analysis of bioinformatics source code. The purpose of our work is to contribute data to the growing conversation in the bioinformatics community around reproducibility, code quality, and software usability. We analyze a large collection of bioinformatics software projects, identifying relationships between code properties, development activity, developer communities, and software impact. Throughout the work, we compare the large set of projects to a small set of highly popular bioinformatics tools, highlighting features associated with high-profile projects. We make our data and code publicly available to enable others to build upon our analysis or generate new datasets. The significance of our work is to (1) contribute a large base of knowledge to the bioinformatics community about the state of their software, (2) contribute tools and resources enabling the community to conduct their own analyses, and (3) demonstrate that it is possible to systematically analyze large volumes of bioinformatics code. This work and the provided resources will enable a more effective, data-driven conversation around software practices in the bioinformatics community.

## Introduction

Bioinformatics is broadly defined as the application of computational techniques to analyze biological data. Modern bioinformatics can trace its origins to the 1960s, when improved access to digital computers coincided with an expanding collection of amino acid sequences and the recognition that macromolecules encode information [1]. The field underwent a transformation with the advent of large-scale DNA sequencing technology and the availability of whole genome sequences such as the draft human genome in 2001 [2]. Since 2001, not only the volume but also the types of available data have expanded dramatically. Today, bioinformaticians routinely incorporate whole genomes or multiple whole genomes, high-throughput DNA and RNA sequencing data, large-scale genetic studies, data addressing macromolecular structure and subcellular organization, and proteomic information [3].

Some debate has centered around the difference between “bioinformatics” and “computational biology”. One common opinion draws a distinction between bioinformatics as tool development and computational biology as science [4]. However, no consensus has been reached, nor is it clear whether one is needed. The terms are often used interchangeably, as in the “Computational biology and bioinformatics” subject area of *Nature* journals, described as “an interdisciplinary field that develops and applies computational methods to analyse large collections of biological data” [5]. In this article we use the umbrella term “bioinformatics” to refer to the development of computational methods and tools to analyze biological data.

In recent years, the explosion of genomic data and bioinformatic tools has been accompanied by a growing conversation around reproducibility of results and usability of software [6–9]. Reproducibility requires that authors publish original data and a clear protocol to allow repetition of the analysis in a paper [7]. Usability refers to ease and transparency of installation and usage. Version control systems such as Git and Subversion, which allow developers to track changes to code and maintain an archive of all old versions, are widely accepted as essential to the effective development of all non-trivial modern software. In particular, transparent version control is important for long-term reproducibility and usability in bioinformatics [6–9].

The dominant version control system today is the open source distributed system Git [10], used by 87.2% of respondents to the 2018 Stack Overflow Developer Survey [11]. A Git “repository” is a directory that has been placed under version control, containing files along with all tracked changes. A “commit” is a snapshot of tracked changes that is preserved in the repository; developers create commits each time they wish to preserve a snapshot. Many online sharing sites host Git repositories, allowing developers to share code publicly and collaborate effectively with team members. GitHub [12] is a tremendously popular hosting service for Git repositories, with 24 million users across 200 countries and 67 million repositories in 2017 [13]. Since its initial launch in 2008, GitHub has grown in popularity within the bioinformatics field, as demonstrated by the proportion of articles in the journal *Bioinformatics* mentioning GitHub in the abstract (Fig 1). For an excellent explanation of Git and GitHub including additional definitions, see [14].

**Fig 1.**
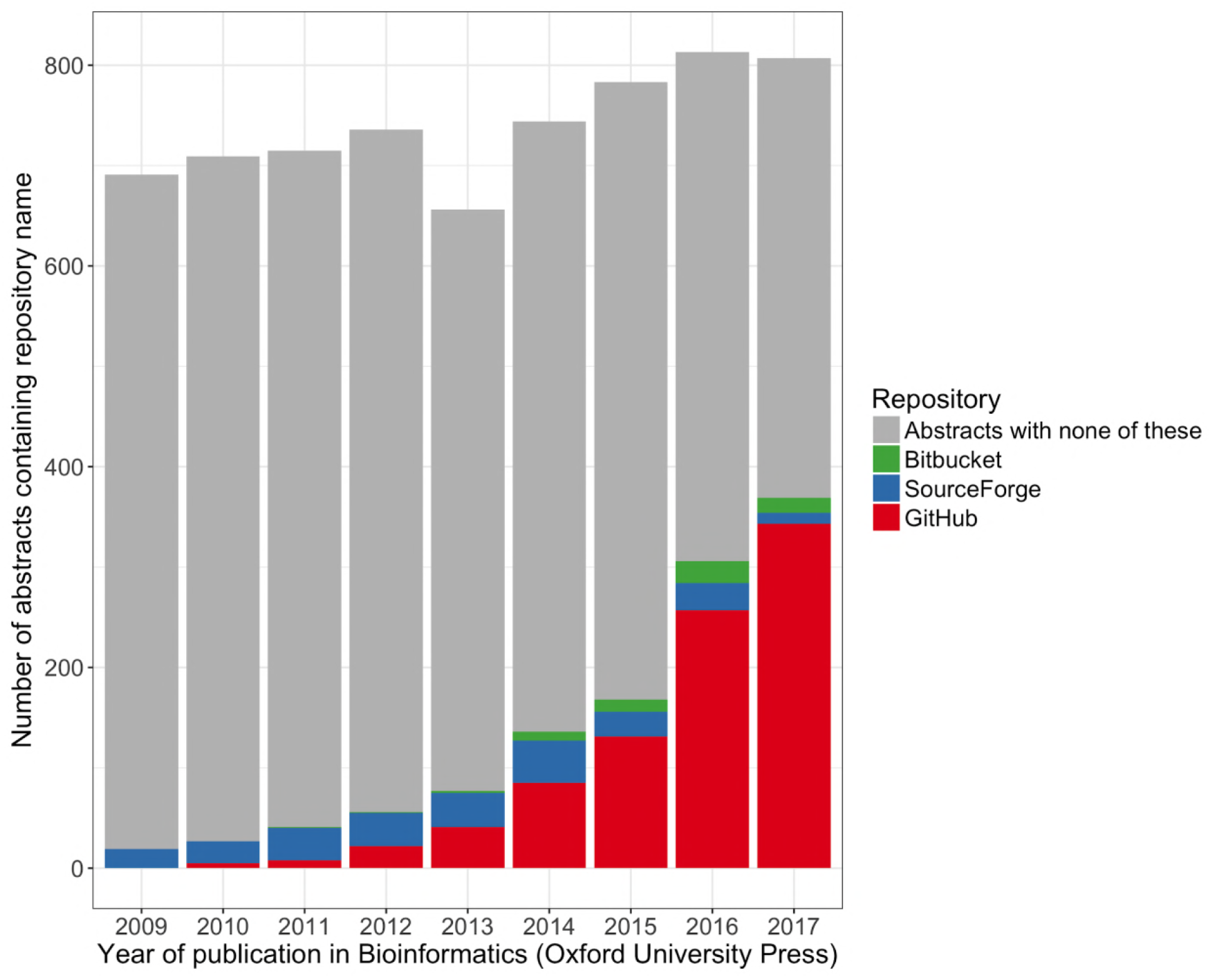
Source code repositories in the journal Bioinformatics. Here the term “repository” refers to online code hosting services. The journal *Bioinformatics* publishes new developments in bioinformatics and computational biology. If a paper focuses on software development, authors are required to state software availability in the abstract, including the complete URL [15]. URLs for software hosted on the popular services GitHub, Bitbucket, and SourceForge contain the respective repository name except in rare cases of developers referring to the repository from a different URL or page. The figure shows the results of PubMed searches for the repository names in the title or abstract of papers published in *Bioinformatics* between 2009 and 2017. The category “Abstracts with none of these” captures all remaining articles published in Bioinformatics for the year, and likely includes many software projects hosted on organization websites or featuring their own domain name, as well as any articles that did not publish software.

The bioinformatics field embraces a culture of sharing — for both data and source code — that supports rapid scientific and technical progress. In this paper, we present, to our knowledge, the first large-scale study of bioinformatics source code, taking advantage of the popularity of code sharing on GitHub. Our analysis data include 1,720 GitHub repositories published along with bioinformatics articles in peer-reviewed journals. Additionally, we have identified 23 “high-profile” GitHub repositories containing source code for popular and highly respected bioinformatic tools. We analyzed repository metadata, source code, development activity, and team dynamics using data made available publicly through the GitHub API [16]. We provide all scripts used to generate the dataset and perform the analysis, along with detailed instructions. We work within the GitHub Terms of Service [17] to make all data except personal identifying information publicly available, and provide instructions to reconstruct the removed columns if needed. Our main analysis results are provided as a table with over 400 calculated features for each repository.

Although the software engineering literature describes many analyses of GitHub data [18–24], bioinformatics software has not been looked at specifically. These software engineering studies often look only at highly active projects in wide community use, with many contributors utilizing the collaborative features of GitHub. Public bioinformatics software serves a variety of purposes, from analysis code supporting scientific results to polished tools intended for adoption by a wide audience. With exceptions, code bases published along with bioinformatics articles tend to be small, with one or a few contributors, and use GitHub mostly for its version control and public sharing features. Additionally, the interdisciplinary nature of bioinformatics creates a unique culture around programming, with developers bringing experience from diverse backgrounds [25]. The projects in our dataset treat a variety of scientific topics, use many different programming languages, and show a diverse range of team dynamics.

We describe our dataset from the perspective of the articles announcing the repositories, the source code itself, and the teams of developers. We observe several features that are associated with overall project impact. Our analysis points to simple recommendations for selecting bioinformatic tools from among the thousands available. Our dataset also contributes to and highlights the importance of the ongoing conversation around reproducibility and software quality.

## Results

### A dataset of 1,740 bioinformatics repositories on GitHub

We curated a set of 1,720 GitHub repositories mentioned in bioinformatics articles in peer-reviewed journals (referred to throughout the paper as the “main” dataset), as well as 23 high-profile repositories that were not necessarily on GitHub at the time of publication or are not published in journals. Three repositories overlapped between the two sets. As a resource for the community, we provide the full pipeline to extract all repository data from the GitHub API, all extracted data except personal identifying information, scripts to perform all analysis, and citations for the articles announcing each repository.

### Article topics

We performed topic modeling [26] on the abstracts of the articles announcing each repository in the main dataset, associating each article with one or more topics. We manually assigned labels to each topic based on top associated terms (Fig S1); for example, the topic “Transcription and RNA-seq” is associated with the terms “rna”, “seq”, and “transcript”. We found that the topic “Web and graphical applications” was positively associated with several measures of project size and activity, as were, to a lesser extent, some other topics (Fig 2). We found that code for articles about certain topics was disproportionately written in certain languages; for example, the greatest amount of code for “Assembly and sequence analysis” was in C and C++, while the greatest amount of code for “Web and graphical applications” was in JavaScript (Fig S2). Bioinformatics was the most common journal for all topics, probably due in part to the relative ease of finding relevant projects in this journal (Fig S3). Fig S4 shows topic distribution by year of initial commit and article publication.

**Fig 2.**
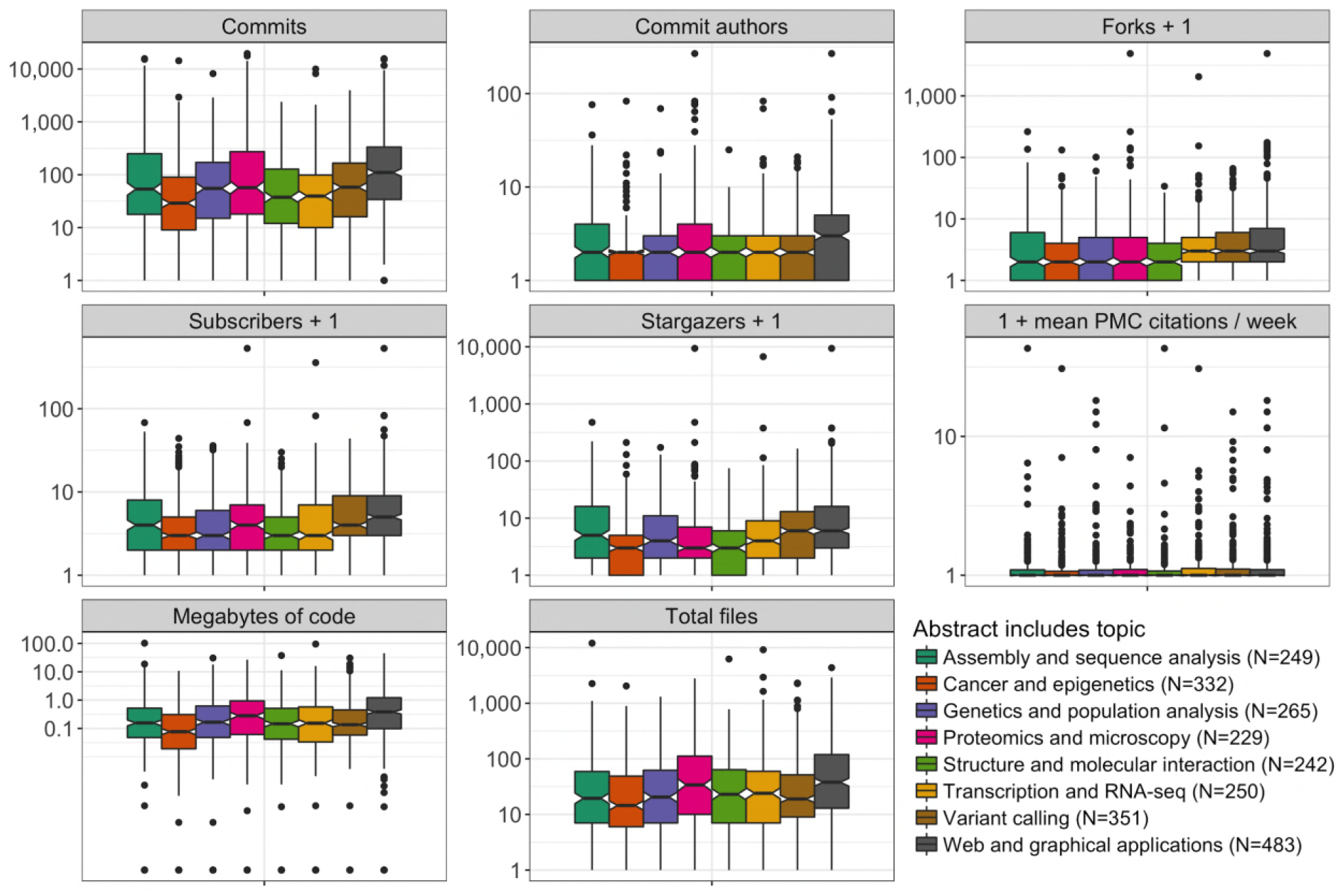
Project features by article topic. Projects are broken into groups according to whether the accompanying paper abstract is associated with each topic category. Projects that are associated with multiple topics are counted separately for each topic. Topic labels were assigned manually after examining top terms associated with each category. We added one to several variables to facilitate plotting on a log scale; these are noted in the variable name. All variables refer to the GitHub repository except “1 + mean PMC citations / week”, which refers to the paper and looks at citations in PubMed Central per week starting two years after the initial publication of the paper. Commits is the total number of commits to the default branch. Commit authors have created commits but do not necessarily have push access to the main branch; we attempted to collapse individuals with multiple aliases. Forks are individual copies of the repository made by community members. Subscribers are users who have chosen to receive notifications about repository activity. Stargazers are users who have bookmarked the repository as interesting. Megabytes of code and total files include source code only, excluding data file types such as JSON and HTML. The horizontal line at the center of the notch corresponds to the median. The lower and upper limits of the colored box correspond to the first and third quartiles. The whiskers extend beyond the hinges by at most an additional 1.5 times the inter-quartile range. Outliers are plotted individually. The notches correspond to roughly a 95% confidence interval for comparing medians [27]. The table of repository features is provided as Table S8.

### Programming languages

We identified a programming language for each source file and analyzed the prevalence of languages along several dimensions including total number of source files, lines of code, and size of source files in bytes. In high-profile repositories, the greatest amount of code in bytes was in Java, followed by C and C++. In the main dataset, two repositories contained entire copies of the large C++ Boost libraries [28]. Ignoring those copies of Boost, the greatest amount of code in the main dataset was in Javascript, followed by Java, Python, C++, and C (Fig S5).

We analyzed language features including primary execution mode (interpreted or compiled), type system (static or dynamic, strong or weak), and type safety. High-profile repositories tended to emphasize compiled, statically typed languages, with the largest contribution being from Java. The main dataset contained a greater proportion of code written in interpreted or hybrid interpreted/compiled (such as Python) and dynamically typed languages (Fig 3, Fig S6, Table S6, Table S7). This difference could reflect the fact that interpreted and dynamically typed languages provide a powerful platform to quickly design prototypes for small projects, while static typing provides important safety checks for larger projects. Indeed, there was a relationship between project size (total lines of code) and amount of statically typed code (percentage of bytes in statically typed languages): the Spearman correlation between these variables over the entire dataset was 0.41 (P= 2.2e-16) (Table S8). Our data support the intuition that Java, Python and R are more succinct than lower-level languages such as C and C++, as the former group tended to have fewer lines of code per source file in the presumably sophisticated high-profile repositories (Fig 3).

**Fig 3.**
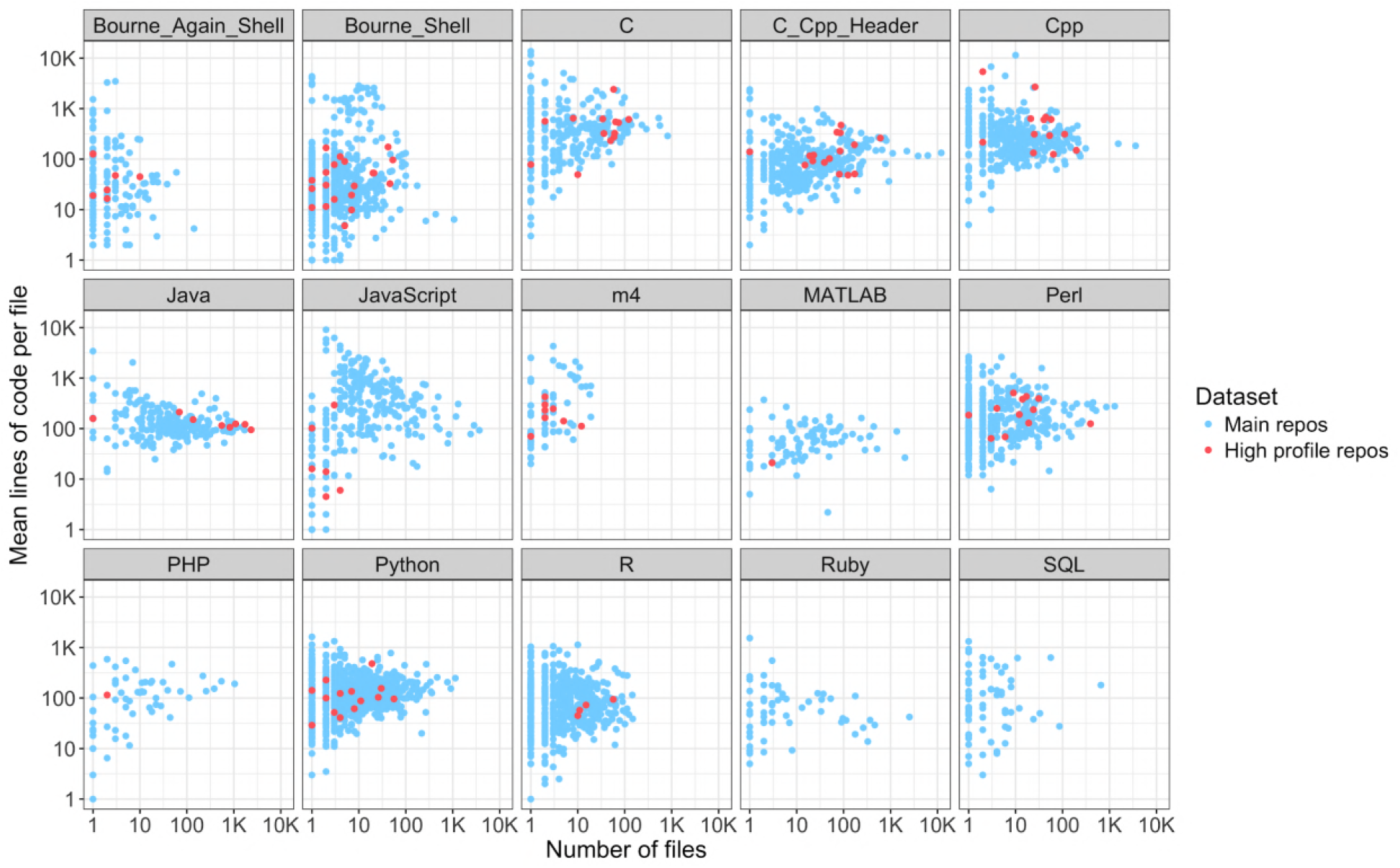
Number and length of source files by programming language. Languages included in at least 50 main repositories are shown. Each dot corresponds to one repository and indicates the number of files in the language and the mean number of lines of code per file not including comments. The data are provided as Table S8.

### Developer communities

For version control systems such as Git, “commits” refer to batches of changes contributed by individual users; each commit causes a snapshot of the repository to be saved along with records of all changes. Each GitHub repository has a core team of developers with commit access; these developers can push changes directly to the repository. In addition, GitHub facilitates community collaboration through a system of forks and pull requests. Anyone can create a personal copy of a public repository, called a “fork”, and make arbitrary changes to their fork. If an outside developer feels their changes could benefit the main project, they can create a “pull request”: a request for members of the core team to review and possibly merge their changes into the main project. In that case, the commit records for the main project would show the outside contributor as the commit author and the core team member who merged the changes as the committer.

We looked at the size of each developer team (including users with commit access and outside contributors) as well as other measures of community engagement, including number of forks, subscribers, and stargazers. Subscribers are users who have chosen to receive notifications about repository activity. Stargazers are users who have bookmarked the repository as interesting. Neither subscribers nor stargazers necessarily touch any code, though in practice they are likely to include the developer team. Not surprisingly, the size of the developer team (all commit authors) was strongly associated with the number of forks, subscribers, and stargazers. High-profile repositories tended to have larger teams and more community engagement by these measures (Fig 4). The number of outside contributors was also associated with these measures, though less strongly, perhaps because only 14% of main repositories had any outside contributors and these already tended to be within the highly active subset; 70% of high-profile repositories had outside contributors (Fig S7).

**Fig 4.**
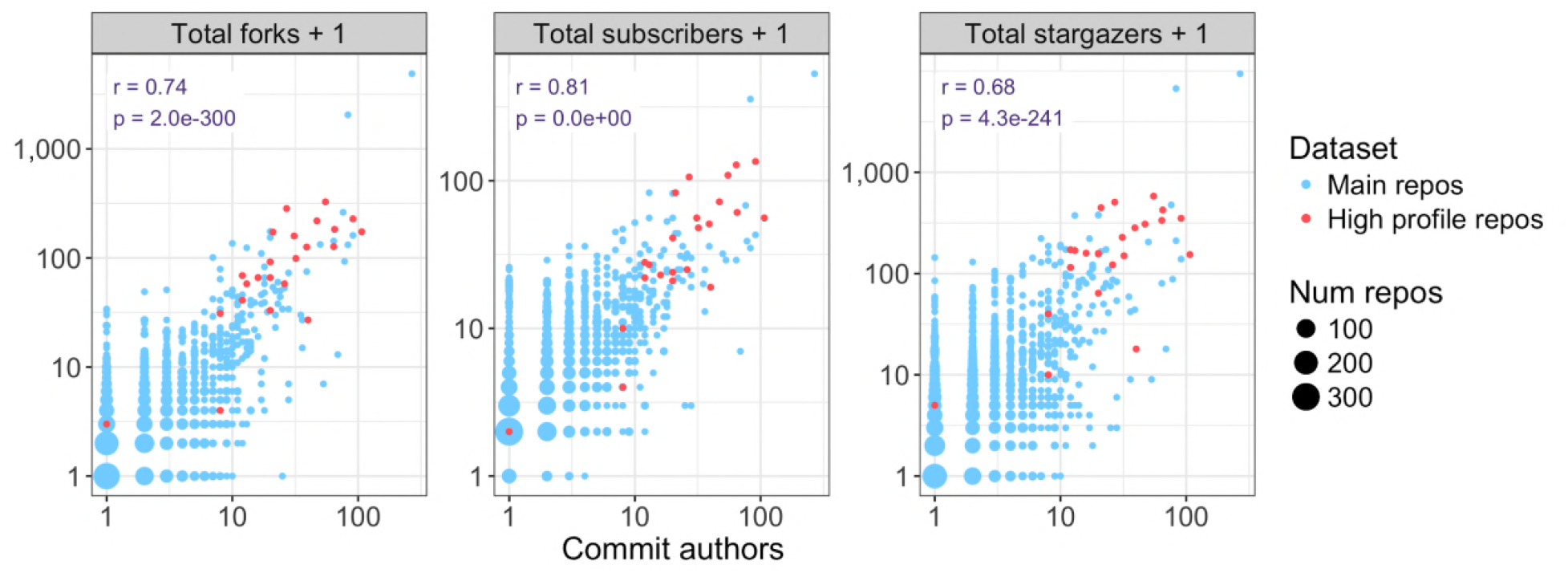
Size of developer community. Various measures of community engagement are plotted against the number of commit authors. Each dot represents one repository or a set of repositories with identical values for the variables. We added one to the vertical axis variables to facilitate plotting on a log scale due to many zero values. The pearson correlation and associated p-value are displayed for each variable versus number of commit authors. Commit authors refers to the number of unique commit authors to the default branch. The high-profile repository with a single contributor is s-andrews/FastQC [29]. This repository appears to have been created by a single developer importing a previously existing code base to GitHub. The table of repository features is provided as Table S8.

### Gender distribution of developers and article authorships

We analyzed the gender distribution of developers and article authorships in the dataset as a whole and within teams. Developer and author first names were submitted to the Genderize.io API [30] and high-confidence gender calls were counted. We found that the proportion of female authors decreased with seniority in author lists and the proportion of female developers was lower in high-profile repositories compared to the main dataset. In the main dataset, 12% of developers were women while only 6% of commits were contributed by women; these numbers were lower in the high-profile dataset (7% and 2%, respectively). In biology articles, it is customary to list the lead author first and the senior author last, with additional authors in the middle. We found that in the articles announcing each repository, middle authors included the greatest proportion of women. Women comprised 22% of all authorships in the main dataset and 21% in the high-profile dataset, compared to 18% and 0% for first authors and 14% and 8% (representing only one person) for the most senior last authors (Fig 5). A separate study of author gender in computational biology articles found a similar trend of decreased representation of women with increased seniority in author lists; the authors additionally identified a pattern of more female authors on papers with a female last author [31].

We analyzed the gender composition of each team of developers and paper authors. The most common type of team in the main dataset was a single male developer and an all-male author list. The most common type of team in the high-profile dataset was a majority-male developer team and an all-male author list. Only ten main repositories and no high-profile repositories had all or majority female developer and author teams; all ten of these developer teams consisted of a single female developer (Fig S8).

We quantified gender diversity within teams using the Shannon index of diversity [32]. A Shannon index of 0 means all members have the same gender, while the maximum value of the Shannon index with two categories is ln(2) = 0.69, achieved with equal representation of both categories. We found that 13% of main repositories and 62% of high-profile repositories had a nonzero Shannon index for the developer team. There were no high-profile repositories with a Shannon index greater than 0.4; the percentage of main repositories with Shannon index greater than 0.4 was 12% (Fig S9).

**Fig 5.**
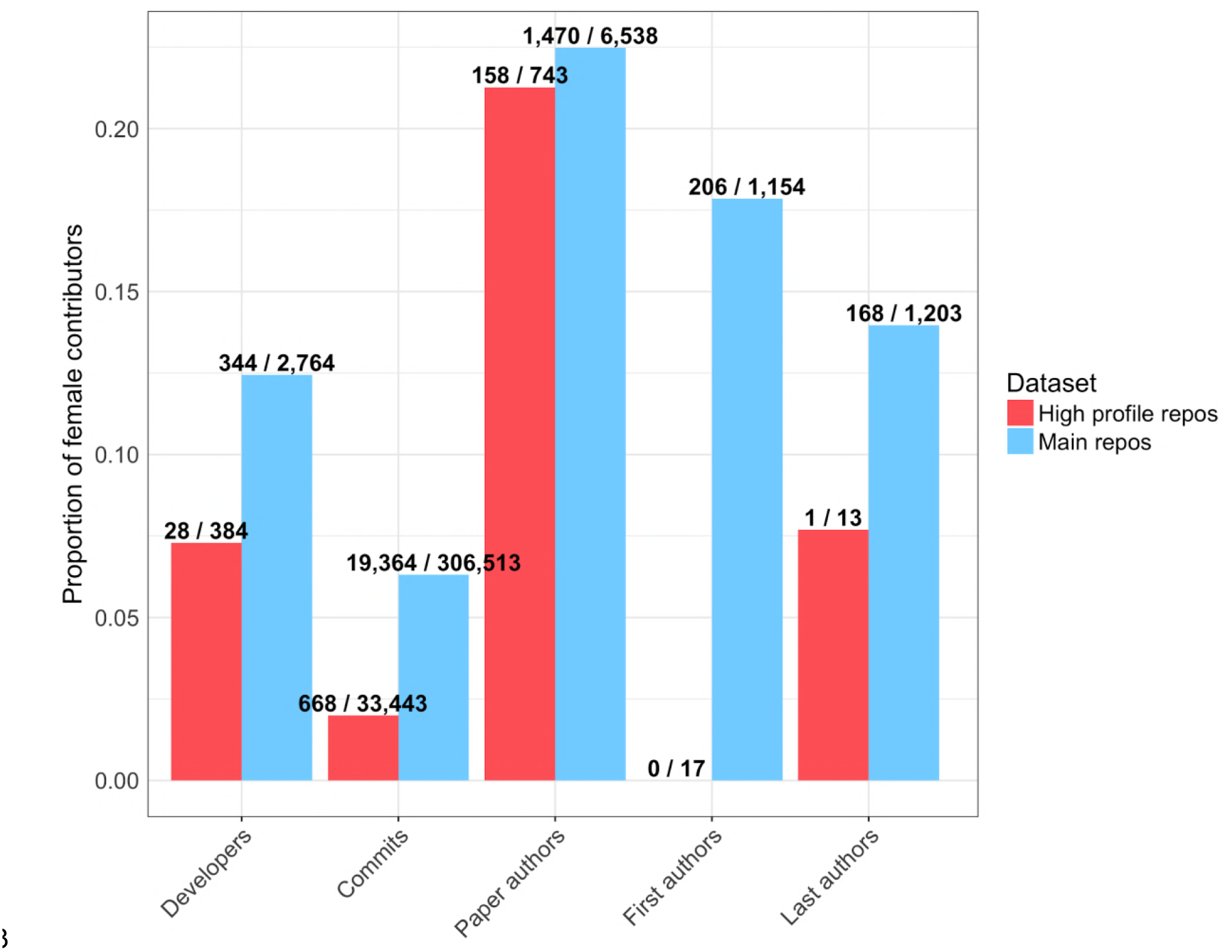
Distribution of developers, commits and paper authorships by gender. “Developers” are unique commit authors or committers over the entire dataset; we attempted to collapse individuals with multiple aliases. “Commits” are individual commits to default branches of repositories. “Paper authors” are individual authorships on papers, not necessarily unique people. For each repository, the one paper announcing the repository is included; papers were then deduplicated because some papers announced multiple repositories. First and last authors are only counted for papers with at least two authors. Names for which a gender could not be inferred are excluded. Bar height corresponds to the number of female contributors divided by the number of contributors with a gender call; these numbers are labeled above each bar. The features for each repo are provided in Table S8.

### Commit dynamics

We looked at several measures of commit timing along with total number of commits to each repository. Not surprisingly, the total number of commits was strongly associated with density of activity (commits per month and maximum consecutive months with commits) and overall project duration. High-profile repositories tended to have longer project duration and greater density of commit activity (Fig 6).

**Fig 6.**
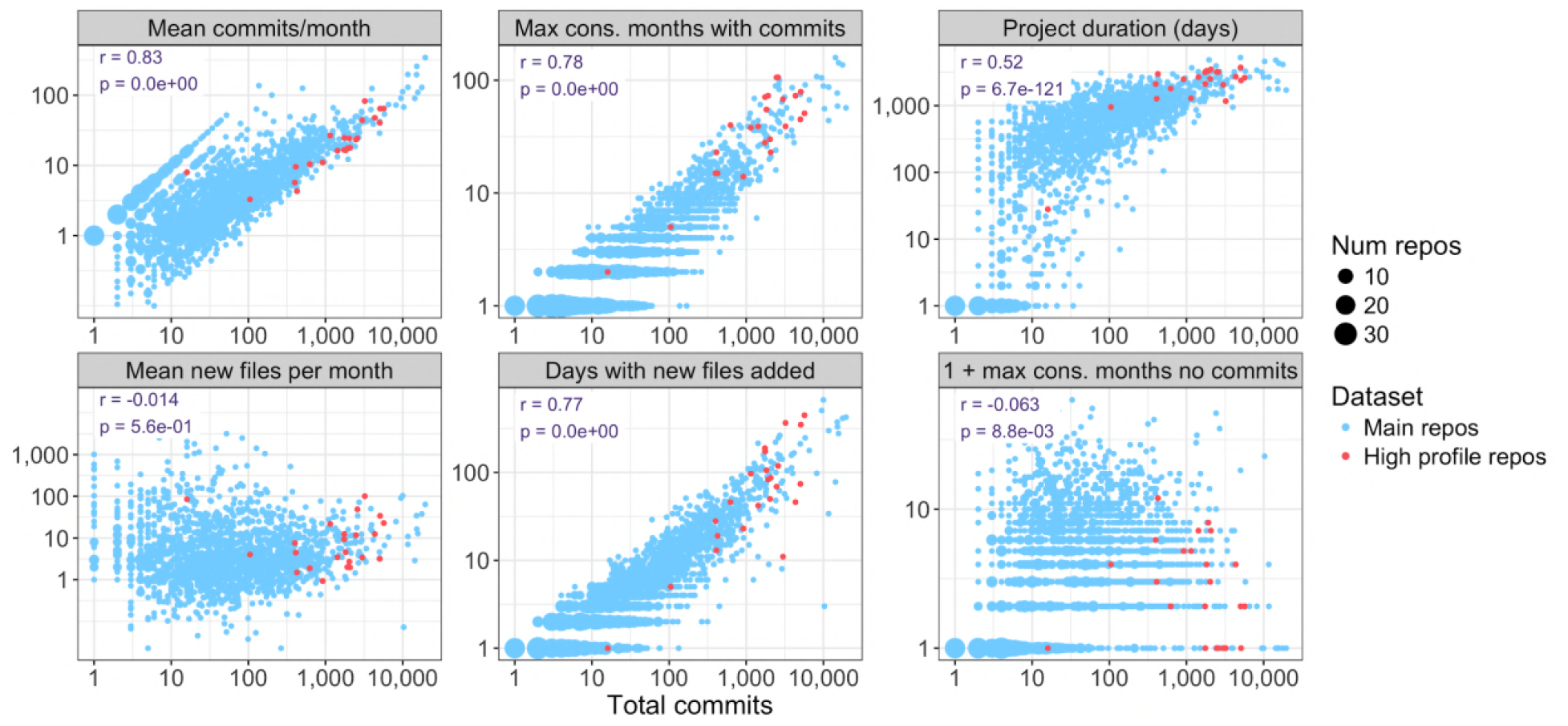
Commit timing versus total commits. Various timing dynamics are plotted versus total commits to the default branch. Each dot represents one repository or a set of repositories with identical values for the variables. For each variable, the total time interval covered by the project is the interval starting with the first commit and ending with the last commit at the time we accessed the data. For example, “Mean new files per month” counts only months from the first to last commit. The high-profile repository with only 16 commits and all files added on a single day is s-andrews/FastQC [29]. This repository appears to have been created by importing a previously existing code base to GitHub. The data are provided as Table S8.

### A simple proxy for project impact

We looked at the simple binary feature of whether any commits were contributed to each repository after the associated article appeared in PubMed. We found that this simple feature was associated with several measures of project activity and impact (Fig 7). Not surprisingly, it was strongly associated with the total number of commits and size of the developer team. Presumably, larger projects tend to be those that are useful to many people and for which development continues after the paper is published. The metric was also associated with measures of community engagement such as forks, stargazers, and outside contributors. This could be explained in part by the previous point and in part by outside community members voluntarily becoming involved in the project after reading the paper. However, interestingly, the association with the proportion of commits contributed by outside authors was not statistically significant, suggesting that overall team size may be the principal feature driving the relationship with the number of outside commit authors. Additionally, the metric was associated with frequency of citations in PubMed Central, which could indicate that people are discovering the code through the paper and using it, and the code is therefore being maintained. Interestingly, repositories with commits after the paper was published had longer commit messages (explanations included by commit authors along with their changes to the repository). This could be due to a relationship between both variables and the size of the developer team; perhaps members of larger teams tend to write longer commit messages to meet the increased burden of communication with more team members. Indeed, there was a moderate linear relationship (r = 0.14, p = 1.9e-09) between total number of commit authors and mean commit message length in the main dataset.

**Fig 7.**
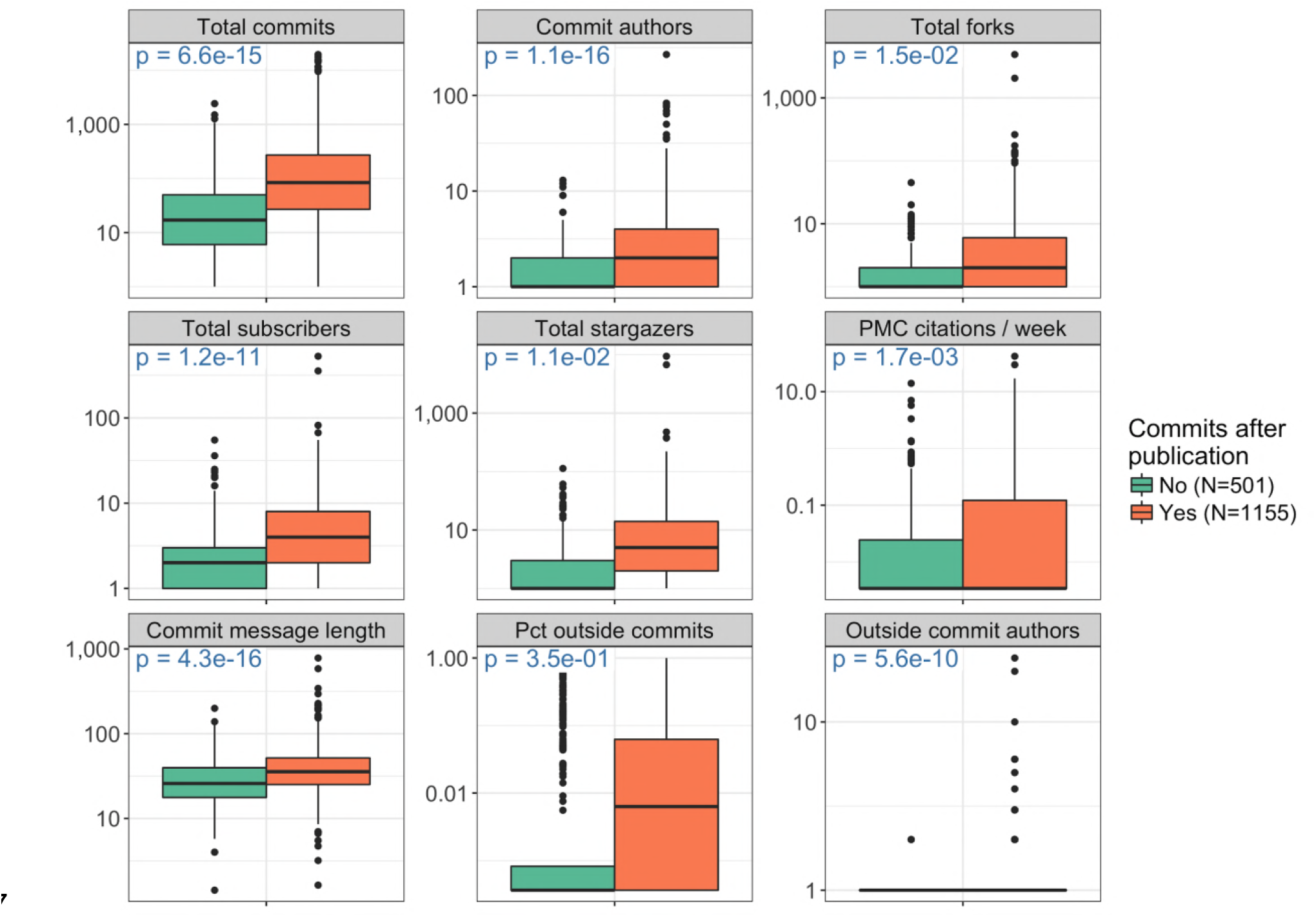
Commits after paper publication. Each data point contributing to each box plot is one repository in the main dataset. Repositories are separated by whether the last commit timestamp at the time we accessed the data was after the date the corresponding publication appeared in PubMed. Repositories for which we do not have a publication date in PubMed are excluded. See Fig 2 legend for the explanation of “Total commits”, “Commit authors”, “Total forks”, “Total subscribers”, “Total stargazers”, and “PMC citations / week”. “Commit message length” is the mean number of characters in a commit message. “Pct outside commits” is the proportion of commits with an author who is never a committer. “Outside commit authors” is the number of commit authors who are never committers. The p-value refers to the two-sided *t*-test for different means between the two groups. The data used to compute the p-value include zero values, but for the plot, we replaced zeros by the minimum positive value of each variable to facilitate plotting on a log scale. The horizontal line across the box corresponds to the median. The lower and upper limits of the box correspond to the first and third quartiles. The whiskers extend beyond the box by at most an additional 1.5 times the inter-quartile range. Outliers are plotted individually. The table of repository features is provided as Table S8.

## Discussion

We have presented the first large-scale analysis of bioinformatics code to our knowledge. Our analysis gives a high-level picture of the current state of software in bioinformatics, summarizing scientific topics, source code features, development practices, community engagement, and team dynamics. The culture of sharing in bioinformatics will continue to enable deeper study of software practices in the field. Our hope is that readers will uncover additional insights in our tables of hundreds of calculated features for each repository (Table S8), many of which were not analyzed in this paper, and that some readers will use or adapt our code to generate data and analyze repositories in unanticipated ways.

Interestingly, despite being made public on GitHub, nearly half of all repositories in our dataset do not feature explicit licenses (Fig S10), in most cases likely unintentionally restricting the rights of others to reuse and modify the code. Nonetheless, the type of research described here may proceed under the GitHub Terms of Service [17] and Privacy Statement [33].

With the overwhelming variety of public bioinformatics software available, users are constantly faced with the question of which tool to use. Several features of our analysis point to simple heuristics based on information available on GitHub. We observed relationships between community engagement and various measures of project size and activity level (Fig 4, Fig 6, Fig S7). Our final analysis looked at the simple question of whether the developers had revisited their code at all after the paper was published; we found that this feature is associated with several measures of impact (Fig 7). Intuitively, these points suggest that users should prioritize software that is being consistently maintained by an active team of developers. The GitHub web interface prominently displays the total number of commits, number of contributors, and time of latest commit on the front page for each repository. Additionally, GitHub provides a full-featured mechanism, called Issues, that allows the developer team or any user to create tracked requests within the project. We did not analyze issues because these are a relatively advanced feature that is rarely used in our dataset; nonetheless, a consistent flow of issues can help identify sophisticated projects under active development.

Bioinformatics is a hybrid discipline combining biology and computer science. There are three major paths into the field: (1) computer scientists and programmers can become familiar with the relevant biology, (2) biologists can learn programming and data analysis, or (3) students can train specifically in increasingly popular bioinformatics programs [25]. Our dataset likely includes developers from all three major paths. However, our analysis of developer gender demonstrates that the gender distribution in bioinformatics more closely resembles that of computer science than biology. Indeed, the underrepresentation of women in our dataset was more extreme than among students awarded PhDs in computer science in the United States in 2016 [34]. A possible reason for this could be that, despite relatively high numbers of women in biology, biologists who make the transition to bioinformatics tend to be male. Another possible explanation could be that the subset of bioinformaticians who publish code on GitHub are disproportionately those from the computer science side. Importantly, our analysis does not address other intersections of identity and demographics that affect individuals’ experience throughout the academic life cycle. Beyond simply pushing for fair treatment of all scientists, researchers have argued that team diversity leads to increased productivity of software development and higher quality science [35–37].

### Limitations

Our dataset represents a large cross section of bioinformatics code bases, but many projects are excluded for various reasons. First of all, due to the challenges of full-text literature search, we did not identify all articles in the biomedical literature that mention GitHub. In particular, we did not use the open access set of articles in PubMed Central because these included too many mentions of GitHub to manually curate for both bioinformatics topics and code being announced with the respective articles, and efforts to train automated classifiers left too many false positives that tended to skew the picture of repository properties compared to true announcements of bioinformatics code. We therefore selected a search strategy that was limited enough to generate a high-quality hand-curated set and could include papers that were not open access. Second, we are missing repositories that were not on GitHub at the time of publication or are primarily described on a main project website other than GitHub, with the exception of the high-profile repositories we added manually. Third, we have not included large open source collaborations such as Bioconductor [38], BioJava [39], and Biopython [40], due to project-specific substructure making it unfair to compare them to the rest of the dataset. Finally, our dataset could be biased due to our use of GitHub itself: it is possible that developers with certain backgrounds are disproportionately likely to host code on GitHub, while we have not analyzed any code not hosted on GitHub.

The spirit of sharing has led to an increase in popularity of preprints: advance versions of articles that have not yet been published in peer-reviewed journals. Preprints can allow scientific progress to continue during the sometimes extensive review process. However, we chose not to include preprints in our literature search for three main reasons. First, we believed that successful peer review was a fair criterion on which to identify serious code bases. Second, we wanted to analyze article metadata that would only be available from databases such as PubMed. Third, the most popular preprint server for biology, bioRxiv [41], does not currently provide an API, putting programmatic access out of reach.

### Future research

Several interesting future analyses are possible with our dataset or extensions to it. First, we did not examine the important topic of software documentation, either within source code or for users. The myriad forms of user documentation (README files, help menus, wikis, web pages, forums, and so on) make this a difficult but important topic to study. Second, static code analysis would provide deep insight into software quality and style. While impractical for a large heterogeneous set of code bases written in many different languages, future studies could uncover valuable insights through focused static analysis of repositories sharing common features. Third, we did not study the behavior of individual developers in depth. Future studies could analyze the social and coding behavior of individuals across all their projects and interests on GitHub. Finally, our analysis does not address the important question of software validity: whether a program correctly implements its stated specification and produces the expected results. The complexity of bioinformatic analysis makes validity testing a very challenging problem. Nevertheless, progress has been made in this area [42–44]. Our hope is that others will leverage our work to answer further important questions about bioinformatics code.

### Toward better bioinformatics software

Our work provides data to enhance the ongoing community-wide conversation around reproducibility and software quality in bioinformatics. Several features of our data suggest a need for community-wide software standards, including the widespread absence of open source licenses (46% of main repositories have no detectable license), the number of repositories not appearing to use version control effectively (12% of main repositories added all new files on a single day, while 40% have a median commit message length less than 20 characters), and the apparent lack of reuse of the software (28% of papers in the main dataset have never been cited by articles in PubMed Central, while 68% have fewer than five citations) (Table S8). Similarly, a study based on text mining found that over 70% of bioinformatics software resources described in PubMed Central were never reused [45]. These orthogonal lines of evidence support the need for the already growing efforts toward supporting better software in bioinformatics and scientific research in general.

Existing efforts to improve research software include the Software Sustainability Institute [46,47], which works toward a mission of improving software to enable more effective research; Better Scientific Software [48], a project that provides resources to improve scientific and engineering software; and Software Carpentry [49–51], which provides highly practical training for research computing. In addition, several reviews recommend specific practices for the software development lifecycle in academic science. In [8], the author provides specific recommendations to improve usability of command line bioinformatics software. The authors of [52] recommend specific software engineering practices for scientific computing. In [9], the authors outline several practices for the entire software development lifecycle. In [53], members of a small biology lab describe their efforts to bring better software development practices to their lab. In [54], the author advocates for changes at the institutional and societal levels that would lead to better software and better science.

Our contribution to this conversation, in addition to the specific conclusions from our analysis, is to demonstrate that it is possible to study bioinformatics software at the atomic level using hard data. With continued updates, this paradigm will enable a more effective, data-driven conversation around software practices in the bioinformatics community.

## Methods

### Identification of bioinformatics repositories on GitHub

GitHub repositories containing bioinformatics code were found through their mention in published journal articles pertaining to bioinformatics topics. Briefly, a literature search identified articles that were likely to pertain to bioinformatics topics and contained mentions of GitHub. Manual curation identified the subset of these articles treating bioinformatics topics, using a detailed definition of bioinformatics. GitHub repository names were automatically extracted from the bioinformatics articles. Mentions of each repository in each article were manually examined to identify repositories containing code for the paper, as opposed to mentions of outside repositories. Repository names were manually deduplicated and fixed for other noticeable issues such as inclusion of extra text due to the automatic parsing of context around the repository name. Repository names were automatically checked for validity using the GitHub API, and repositories with issues in this check were manually fixed or removed if the repository no longer existed. The final set included 1,720 repositories. In addition to the 1,720 repositories identified through the literature search, we also curated a separate set of 23 high-profile repositories — highly popular and respected tools in the bioinformatics community — based on the high volume of posts about these projects on the online forum Biostars [55]. The two datasets are referred to throughout the paper as the “main” and “high-profile” datasets. See Supplemental Section 2 for details. The repositories are listed in Table S4 and Table S5.

### Extraction of repository data from GitHub API

Repository data were extracted from the GitHub REST API v3 [16] and saved to tables on Google BigQuery [56] for efficient downstream analysis. Data extracted for each repository include repository-level metrics, file information, file creation dates, file contents, commits, and licenses. GitHub API responses were obtained using the PycURL library [57]. The JSON responses were converted to database records and pushed to tables on BigQuery using the BigQuery-Python library [58]. See Supplemental Section 3 for details.

### Topic modeling of article abstracts

We used latent Dirichlet allocation (LDA) [59] to infer topics for abstracts of the articles announcing each repository in the main dataset. From the LDA model, we identified terms that were primarily associated with a single topic. We chose a model with eight topics due to its maximal coherence of concepts within the top topic-specialized terms. We manually assigned a label to each of the eight topics that captures a summary of the top terms. We then classified each article abstract into one or more topics. Details are in Supplemental Section 4.

### Programming languages

We identified 515,017 total files files among the repositories in the main dataset and 22,396 total files in the high-profile dataset. Contents of 425,967 and 18,501 files respectively (349,834 and 16,917 with unique contents) with size under 999KB were saved to tables in BigQuery for further analysis. (See Supplemental Section 3.) We used cloc (Count Lines of Code) version 1.72 [60] to identify the programming language, count lines of code and comments, and extract comment-stripped source code for each file. A total of 221,343 unique files in the main dataset and 11,425 in the high-profile dataset had an identifiable programming language. Language execution modes were obtained from [61]. Type systems were obtained from [62]. Further details are presented in Supplemental Section 5.

### Developer communities

We identified the number of commit authors and outside contributors for each repository. For commit authors, we attempted to count unique people by collapsing users with the same name or login. For outside contributors, we counted commit authors whose author ID is never a committer ID for the repository. The counts of forks, subscribers and stargazers were returned directly from the GitHub API. Further details are presented in Supplemental Section 6.

### Gender analysis

We attempted to infer a gender for each commit author, committer, and article author using the Genderize.io API [30], which returns a gender call and probability of correctness for a given first name. Names were first cleaned to remove noise such as single-word handles or organization names, and then the first word of each cleaned full name was submitted to Genderize. We accepted gender calls whose reported probability was 0.8 or greater. We proceeded with analysis of “female” and “male” categories only. We assume that transgender and non-binary contributors have names that reflect their gender identity. There may be erroneous calls for individuals who do not identify with a binary gender. The gender calls are also expected to include a few errors for cisgender individuals as we accept calls with global probability of 0.8 or higher.

To analyze the gender breakdown of developers, we counted unique full names of authors and committers. For commits, we joined commit records to genders by the full name of the commit author and counted individual commits. For paper authors, we counted individual authorships on papers instead of unique individuals, reasoning that multiple different authorships for the same individual should be counted separately. We analyzed team composition for the 504 projects in the main dataset for which we could infer a gender for at least 75% of developers and 75% of paper authors (Fig S8). We calculated the Shannon index of diversity [32] for the 602 repositories in the main dataset for which we could infer a gender for at least 75% of developers (Fig S9). Details are described in Supplemental Section 7.

### Commit dynamics

We defined project duration as the time span between the first and last commit timestamps for the repository. Metrics describing monthly activity are with respect to the number of months in the project duration. We identified the initial commit time for each file by taking the earliest timestamp of all commits touching the file. Details are described in Supplemental Section 8.

### Proxy for project impact

We defined “commits after publication” to be true if the latest commit timestamp at the time we accessed the data was after the day the associated article appeared in PubMed. Articles were identified and article metadata were extracted as described in Supplemental Section 2.

Repository data were extracted from the GitHub API as described in Supplemental Section 3. Details are described in Supplemental Section 9.

### Availability of data and software

All repository data extracted from the GitHub API, except file contents, are available at https://doi.org/10.17605/OSF.IO/UWHX8. For file contents, in the absence of explicit open source licenses for the majority of repositories studied, we recorded the Git URL for the specific version of each file so that the exact dataset can be reconstructed using our downstream scripts. Additionally, we have removed personal identifying information from commit records, but have included API references for each commit record so that the full records can be reconstructed. Software to generate the dataset and replicate the results in the paper is available at https://github.com/pamelarussell/github-bioinformatics. See Supplemental Section 1 for details on the data and software.

## Acknowledgements

We thank Debashis Ghosh, Wladimir Labeikovsky, and Matthew Mulvahill for helpful conversations and comments on the manuscript. We thank the GitHub support staff for their effort in determining how we could work within the GitHub Terms of Service to publish a reproducible study.

## Author contributions

PR: Conceptualization, Data Curation, Formal Analysis, Methodology, Project Administration, Resources, Software, Supervision, Visualization, Writing – Original Draft Preparation, Writing – Review & Editing.

RJ: Data Curation, Writing – Review & Editing.

SA: Conceptualization, Writing – Review & Editing.

BH: Data Curation, Investigation, Writing – Original Draft Preparation, Writing – Review & Editing.

NC: Funding Acquisition, Project Administration, Supervision, Writing – Review & Editing.

## Supporting information captions

**Supplemental Information. Supplemental information, methods, and figures**.

**Table S1. Definition of bioinformatics topics.**

**Table S2. Manual classification of articles as bioinformatics or not.**

**Table S3. Automatic identification of GitHub repository names in articles.**

**Table S4. Manual curation of GitHub repository names.**

**Table S5. High-profile repositories.**

**Table S6. Programming language type systems.**

**Table S7. Programming language execution modes.**

**Table S8. Calculated repository features.**

